# Ex vivo model of breast cancer cell invasion in live lymph node tissue

**DOI:** 10.1101/2024.07.18.601753

**Authors:** Katerina Morgaenko, Abhinav Arneja, Alexander G. Ball, Audrey M. Putelo, Jennifer M. Munson, Melanie R. Rutkowski, Rebecca R. Pompano

## Abstract

Lymph nodes (LNs) are common sites of metastatic invasion in breast cancer, often preceding spread to distant organs and serving as key indicators of clinical disease progression. However, the mechanisms of cancer cell invasion into LNs are not well understood. Existing in vivo models struggle to isolate the specific impacts of the tumor-draining lymph node (TDLN) milieu on cancer cell invasion due to the co-evolving relationship between TDLNs and the upstream tumor. To address these limitations, we used live ex vivo LN tissue slices with intact chemotactic function to model cancer cell spread within a spatially organized microenvironment. After showing that BRPKp110 breast cancer cells were chemoattracted to factors secreted by naïve LN tissue in a 3D migration assay, we demonstrated that ex vivo LN slices could support cancer cell seeding, invasion, and spread. This novel approach revealed dynamic, preferential cancer cell invasion within specific anatomical regions of LNs, particularly the subcapsular sinus (SCS) and cortex, as well as chemokine-rich domains of immobilized CXCL13 and CCL1. While CXCR5 was necessary for a portion of BRPKp110 invasion into naïve LNs, disruption of CXCR5/CXCL13 signaling alone was insufficient to prevent invasion towards CXCL13-rich domains. Finally, we extended this system to pre-metastatic TDLNs, where the ex vivo model predicted a lower invasion of cancer cells. The reduced invasion was not due to diminished chemokine secretion, but it correlated with elevated intranodal IL-21. In summary, this innovative ex vivo model of cancer cell spread in live LN slices provides a platform to investigate cancer invasion within the intricate tissue microenvironment, supporting time-course analysis and parallel read-outs. We anticipate that this system will enable further research into cancer-immune interactions and allow isolation of specific factors that make TDLNs resistant to cancer cell invasion, which are challenging to dissect in vivo.

## INTRODUCTION

Breast cancer is one of the most common primary cancers worldwide, annually diagnosed in > 270,000 patients.^1^ In breast cancer, metastatic disease remains the underlying cause of mortality,^2^ and it occurs preferentially through the lymphatics, with 8-fold higher invasion of lymphatics than blood vessels.^3^ The sentinel lymph node (LN), located downstream from the primary cancer, is the first organ contacted by cancer cells passing through the lymphatic vessels and may provide a niche for metastatic seeding.^4^ Indeed, 27% of breast cancer patients have detectable LN metastasis at diagnosis.^5^ The presence of LN metastasis is linked to poorer survival outcomes compared to patients without nodal involvement,^6^ potentially due to induction of immune tolerance^7^ and/or subsequent dissemination to distant organs.^8–10^ However, despite its potential importance to patient outcomes, the factors fostering a favorable milieu for cancer cell infiltration of the LN and the underlying mechanisms governing this process remain incompletely understood.

Cancer cells that reach the TDLN encounter a highly organized lymphoid structure in the midst of change. Designed for survey of antigens draining from upstream organs, the LN can be compartmentalized into four major anatomical regions: subcapsular sinus (SCS), B cell-rich cortex, the T cell-rich paracortex, and the medulla (Figure 1A). Before metastatic seeding occurs, TDLNs undergo extensive structural and functional remodeling.^11^ Structurally, lymphangiogenesis and enlargement of high endothelial venules,^12^ dilation of the SCS,^13^ and a relaxation of the underlying stromal network collectively affect size exclusion^14^ and fluid permissiveness^15^ of lymphatic conduits. Furthermore, the secretion of chemokines in TDLNs dynamically changes in response to the upstream tumor.^11^ However, little is known about how all of these changes cumulatively impact the receptivity of the TDLN to cancer cell invasion. Some evidence suggests that the tumor primes its TDLN to be more receptive to metastasis than non-draining LN,^16–18^ while other evidence indicates that tumor-induced remodeling of TDLN facilitates immune priming and elimination of cancer at early stages.^19–21^

**Figure 1.**
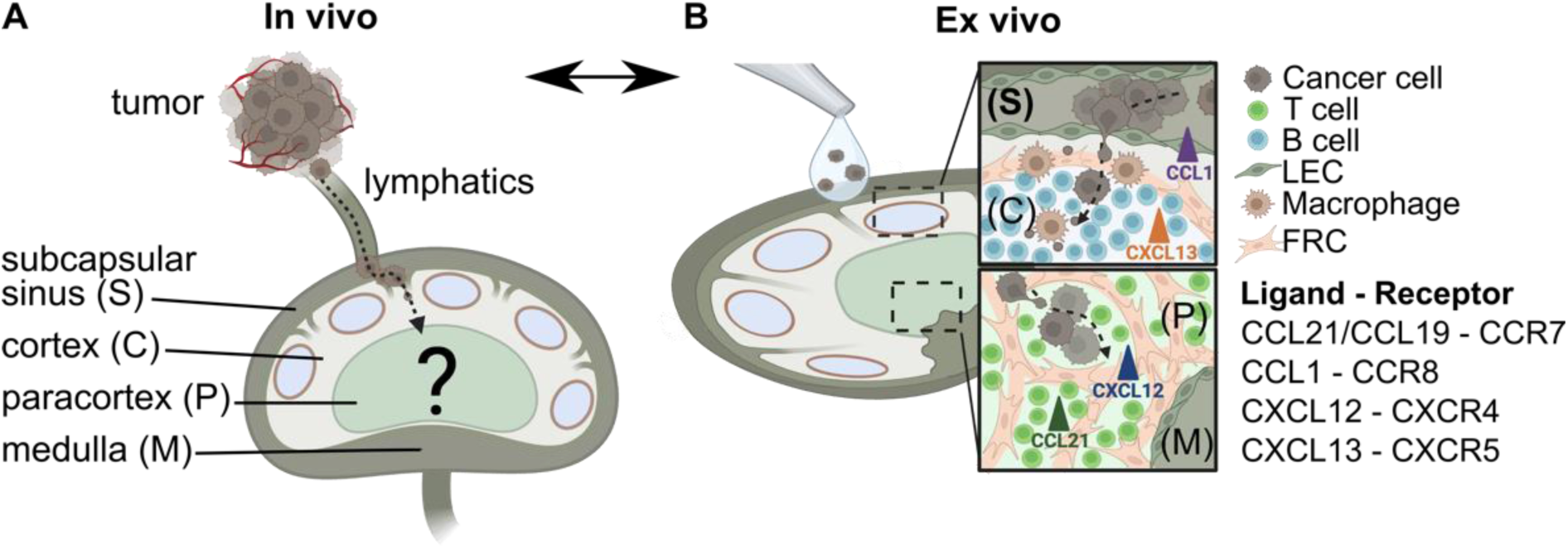
Conceptual illustration of an ex vivo model using live LN tissue slices to model cancer cell chemotaxis in TDLNs. (A) In vivo, cancer cells from the primary tumor invade the lymphatic system and eventually the TDLN, where mechanisms of invasion are difficult to parse. Anatomical zones of LNs include subcapsular sinus (S), cortex (C), paracortex (P) and medulla (M). (B) An ex vivo model of chemotactic invasion of cancer cells within the organized LN architecture. Insets show spread of cancer cells in distinct anatomical regions of the LN. List of chemokine ligand – receptor signaling axes considered in this work. Figure created with BioRender.com.

Locations of invasion and survival in the LN are likely influenced by local microenvironmental cues such as chemokines and cellular activity. Cancer cells often enter the TDLN through the SCS and then penetrate deeper into the cortex via the lymphatic barrier at the sinus floor.^13^ There is strong evidence that chemokines facilitate cancer cell migration from the tumor site into the lymphatics and TDLN, with cancer cells often exploiting the same homing mechanisms used by leukocytes to reach specific regions of LN.^4^ However, many questions remain, including which regions of the LN preferentially support invasion, to what extent cancer cells invade chemokine-rich domains, whether blockade of chemokine signaling could modulate LN metastasis, and even whether the pre-metastatic TDLN is primed to be more or less receptive to invasion.

Questions such as these are challenging to answer using existing models, especially when accounting for the dynamic state of the LN. Most studies are performed in vivo in animal models, and these systems significantly improved our understanding of cancer cell metastasis in TDLN. However, the TDLN co-evolves with the tumor in vivo, making it difficult to study how invasion behavior may depend on the state of the LN separately from how it depends on the tumor microenvironment. In vivo, it is hard to discern how drugs or gene modifications made to the cancer cells may separately impact egress from the primary tumor, entry into primary lymphatics, and invasion into the LN itself. Furthermore, assessing the dynamics of cancer cell invasion within specific LN regions over time is technically challenging, due to the terminal nature of most imaging approaches, limited numbers of reporter animal models, and the complexity of advanced in vivo imaging.^22,23^ For these reasons, a variety of 3D cell culture systems have been developed to recapitulate features of LN architecture and signaling cues in the context of cancer metastasis. These systems have mimicked the microenvironment or fluid dynamics of specific anatomical regions of TDLNs;^24,25^ recreated molecular communication between immune and tumor compartments;^26^ and allowed for the testing of the effects of microenvironmental cues and immunotherapies on tumor cell survival.^27–29^ While these systems potentially enable precise control of the microenvironment and allow time-course analysis, to date no model has captured the dynamic events of cancer cell invasion and spread in the spatially organized LN, nor replicated the role of chemokine signaling in cancer cell invasion of the LN parenchyma.

More than three decades ago, Brodt pioneered the use of frozen murine LN sections and demonstrated a correlation between cancer cell attachment to the 2-dimensional LN sections in vitro and their potential for lymphatic metastasis in vivo.^30^ Recent work has shown that live LN explants support 3D cell migration and spread through organized tissue and maintain chemotactic function.^31–33^ However, although T cell motility is commonly studied in LN slices,^31,34^ cancer cell invasion has not been tested.

Here we aimed to establish a new ex vivo model on LN metastasis based on live ex vivo LN slices (Figure 1B). We tested the hypothesis that the chemotactic activity in live LN slices could recruit cancer cells into the LN parenchyma and predict aspects of the dynamic distribution of cancer cells previously reported in vivo. We tested the extent to which invasion was driven towards particular chemokines, and demonstrated how the model could be used to test requirements for chemokine signaling in cancer invasion. Finally, we applied this system to model invasion into pre-metastatic TDLNs, to begin to address an open question of whether pre-metastatic nodes are more permissive or resistant to invasion.

## RESULTS AND DISCUSSION

### BRPKp110 breast cancer cells were chemoattracted to chemokines secreted by live naïve LN tissue slices

Approximately 75% of breast carcinomas fall into the category of hormone receptor-positive (HR+) due to the expression of estrogen receptor and/or progesterone receptor.^35^ Therefore, for this study, we selected a HR+ murine mammary cancer cell line, BRPKp110. BRPKp110 was established by culture of primary mammary carcinomas after p53 ablation and the transgenic expression of an oncogenic form of K-ras, which is commonly found in human breast cancers.^31^ Similar to human breast cancer carcinomas, in vivo inoculation of BRPKp110 into immune competent mice leads to lymphovascular invasion into TDLNs, making it a good choice to model LN metastasis.^36^

As a first step towards establishing an ex vivo model, we assessed the ability of breast cancer cells to migrate towards conditioned media (CM) from LN slice cultures in vitro. In a 3D transwell assay (Figure 2A), CM from overnight culture of naïve murine LN tissue slices promoted a significant increase in BRPKp110 migration in comparison to control media (Figures 2B, C). This effect was abolished in cancer cells pretreated with Pertussis toxin (PTx), suggesting migration was mediated via chemokine signaling. To rule out potential off-target effects, we verified that PTx treatment did not alter BRPKp110 actin morphology nor affect proliferation rate (Figure S1A).

**Figure 2.**
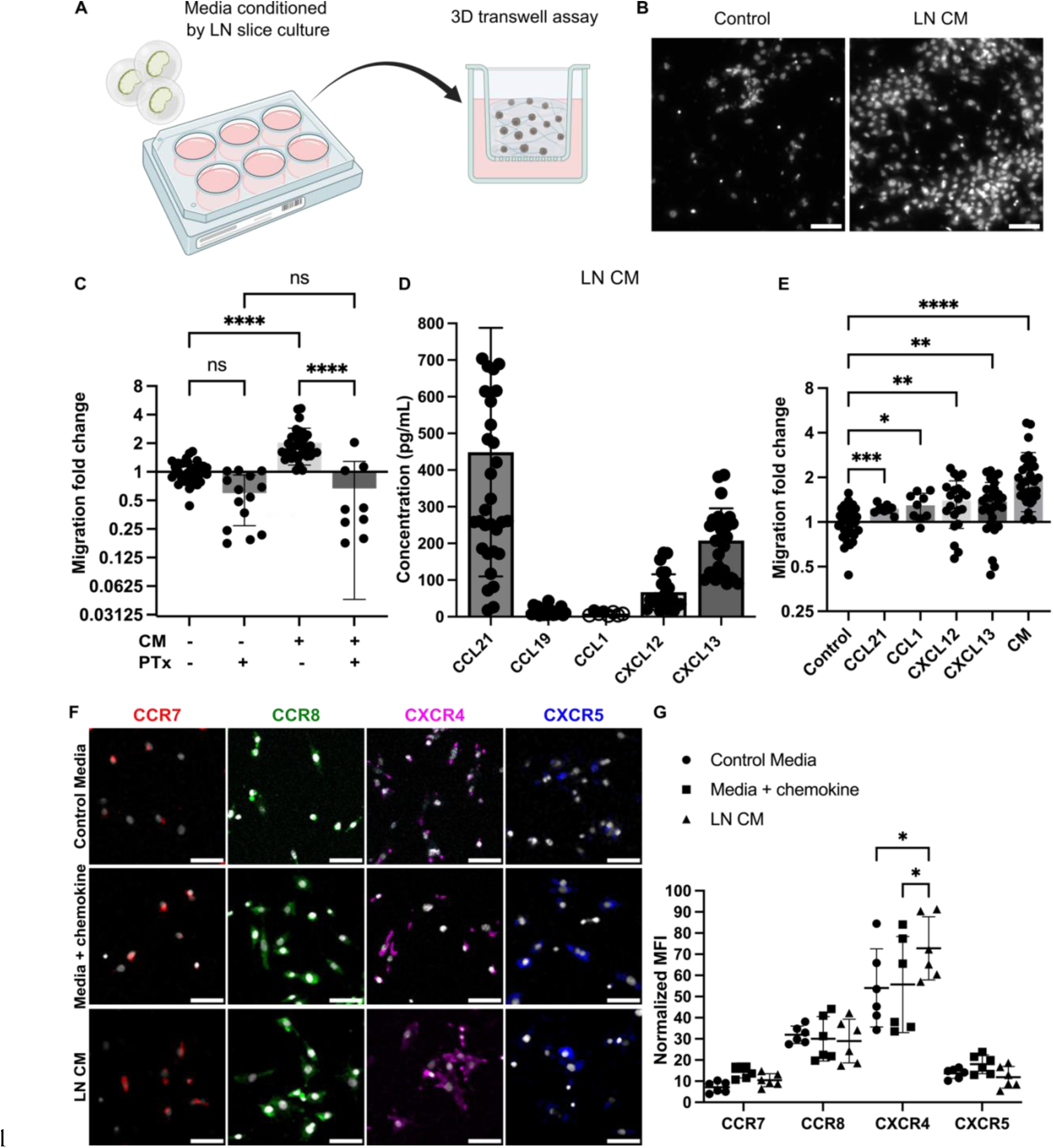
**Naïve LN CM promotes chemotactic migration of BRPKp110 breast cancer cells**. (A) Experimental schematic of 3D transwell migration assay, in which cancer cells in hydrogel were added to the upper compartment and allowed to migrate overnight towards control or conditioned media in the lower compartment. (B) Representative images of the invasion of BRPKp110 cells through the transwell membrane towards control media and media conditioned by LN slice culture. Scale bar 100 µm. (C) Migration data towards conditioned media, normalized to the mean of the migration towards control media. Mean ± stdev; each data point represents one membrane (n=3-5/condition; pooled from 3 independent experiments). Two-way ANOVA with Sidak posthoc test. ****p &lt; 0.0001. (D) Concentrations of CCL21, CCL19, CCL1, CXCL12 and CXCL13 in CM from LN slices after 20 hr culture, measured by ELISA. Mean ± stdev; each dot shows the supernatant from one LN slice. n = 15-35 slices, pooled from 5 female mice. An unfilled circle indicates measurement below the limit of detection. (E) Migration data towards media supplemented with individual chemokines at a concentration of 100 ng/mL, and towards CM, normalized to the mean of the migration towards control media. Mean ± stdev; each data point represents migration fold change per membrane (n=3-5/group; normalized data pooled from 3 independent in vitro experiments). One-way ANOVA, with Dunnett posthoc test. *p &lt;0.05, **p &lt; 0.01, ***p = 0.001, ****p &lt; 0.0001. (F) Representative images of surface immunofluorescence of chemokine receptors on BRPKp110 breast cancer cells after culture in control media, media supplied with the respective chemokine at 200 ng/mL, or LN CM. Scale bar 100 µm. (G) Quantification of receptor expression under various culture conditions. MFI of chemokine receptors across the image was normalized to cell count. Mean ± stdev; each data point represents the average across one culture well; data pooled from 3 independent experiments of 2 replicate wells. Two-way ANOVA with Tukey posthoc test. *p &lt;0.05.

Next, we sought to identify the chemotactic stimuli secreted by live naïve LN slices. Clinical research has shown correlations between CCL21, CCL19/CCR7, CXCL12/CXCR4 and CXCL13/CXCR5 signaling and extensive lymphatic spread and increased risk of LN metastasis in breast cancer^38–44^ and pancreatic ductal adenocarcinoma.^45,46^ The CCL21/CCR7 axis also promoted migration of metastatic melanoma cells towards lymphatics in vitro and in vivo,^47,48^ and the CCL1/CCR8 axis control cancer cell entry into the sinus of the TDLNs in vivo.^13^ Therefore, we measured the levels of these chemokines in the CM. In overnight culture, live LN tissue slices secreted detectable levels of CCL21, CCL19, CXCL12 and CXCL13, whereas CCL1 was below the level of detection (Figure 2D). Media supplemented with recombinant versions of these chemokines individually resulted in an increase in cancer cell migration, but to a lesser extent than towards CM (Figure 2E), suggesting that some synergy may occur towards the mixture of chemokines present in the CM.

Because chemokine signaling requires receptor expression on the cancer cells, we next tested chemokine receptor expression on BRPKp110 cells. Immunofluorescence labeling indicated that BRPKp110 cells expressed all four of the cognate surface receptors: CCR7, CCR8, CXCR4 and CXCR5 (Figure 2F; unstained controls shown in S1B). Interestingly, CXCR4 expression was notably increased in cells cultured in LN CM compared to in control media or media supplemented with CXCL12 (Figures 2G), suggesting regulation by LN-secreted signals. BRPKp110 cells also responded to the CM and to individual chemokines with cytoskeletal rearrangements (F-actin staining) and altered cell morphology from elongated to round (Figure S1C), further confirming their responsiveness to these ligands.

Collectively, these data demonstrated that BRPKp110 cells were chemoattracted to chemokines secreted by LN tissue and expressed functional receptors for the relevant chemokines, suggesting the potential for chemotactic migration into LN tissue.

### Cancer cells infiltrated and proliferated in live ex vivo LN slices

To move from culture inserts to invasion into structured tissue, we tested the extent to which ex vivo LN slices could support cancer cell seeding, invasion, and spread. We developed a procedure in which a suspension of fluorescently labelled, syngeneic BRPKp110 cells was seeded on top of 300-μm thick live LN slices from naïve C57BL/6J female mice, incubated for 1 hr, and washed to remove excess cells (Figure 3A). We found that many cells were washed away, so that only a fraction of overlaid cells had penetrated into the tissue. We refer to this procedure as an “overlay” of cancer cells onto the tissue slices. After the overlay, the tissues were labelled via live immunofluorescence to identify LN zones.^32^ In preliminary work, we determined an optimal seeding density of 20,000 cancer cells per LN slice by seeding various densities onto LN slices (data not shown).

**Figure 3.**
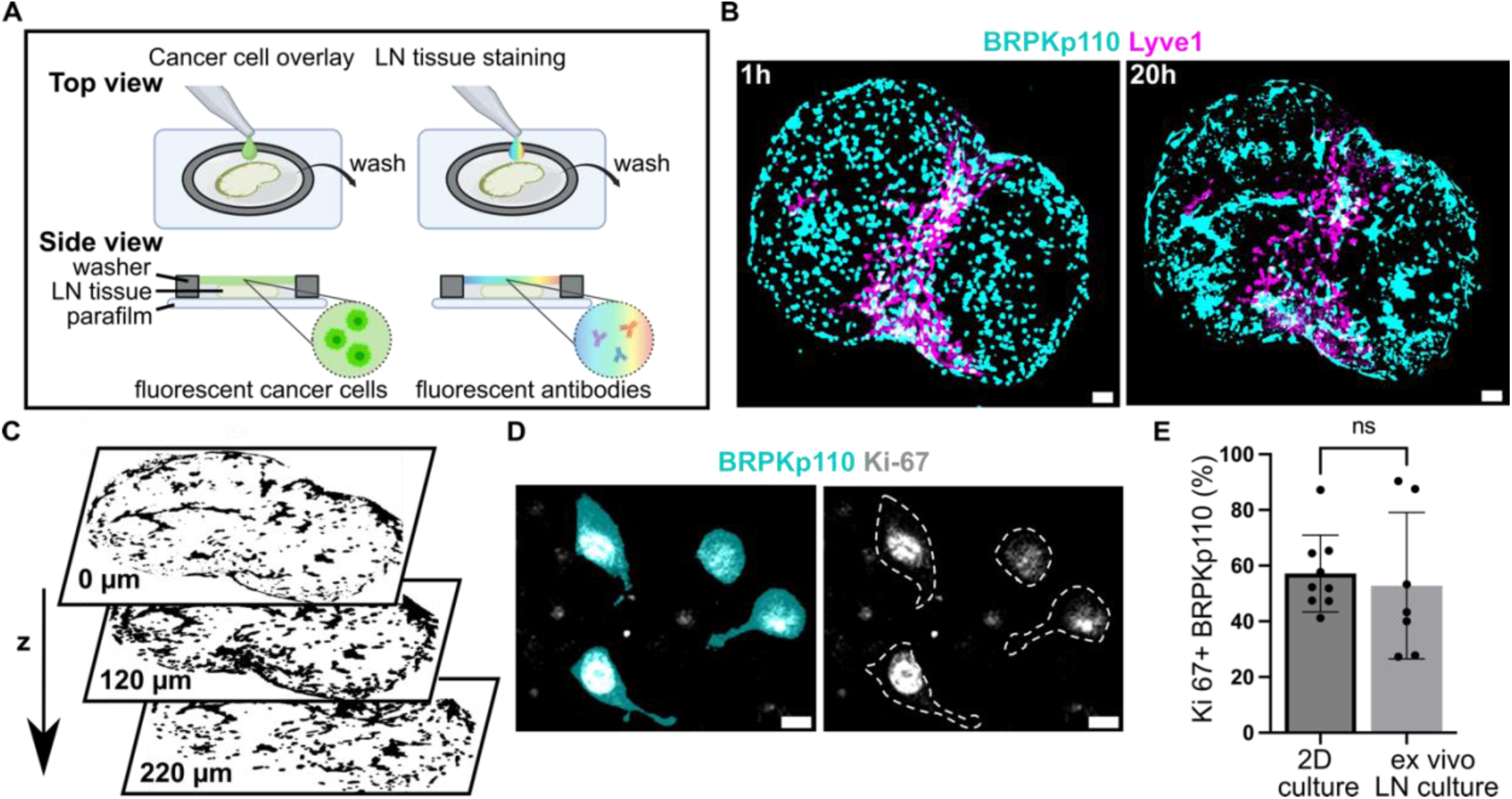
Cancer cells introduced to live LN slices ex vivo infiltrate, proliferate and exhibit a dynamic spreading over a 20 hr culture period. (A) Schematic representation of cancer cell seeding onto live 300-μm sections of LN tissue, followed by live immunostaining via fluorescently conjugated antibodies. (B) Fluorescent BRPKp110 cells (NHS-Rhodamine, cyan) were seeded ex vivo onto naïve LN slices stained for lymphatic endothelial cells (lyve1, magenta) and imaged at 1 hr and 20 hr after seeding. Scale bar 200 µm. (C) Binary image of cancer cells at multiple z-depths illustrating infiltration into the LN tissue. (D) Representative image of proliferating BRPKp110 cells (NHS-Rhodamine, cyan) positive for Ki-67 (gray) 20 hr after seeding onto LN tissue. The left image shows merged channels for BRPKp110 and Ki-67; the right image displays Ki-67 with cell contours outlined by a dotted line. Scale bar 20 µm. (E) Percent of Ki-67 positive cells per field of view in BRPKp110 cultured for 20h alone or seeded ex vivo onto live LN. Mean ± stdev; each data point represent measurement from an individual sample (n=2-3/group, data pooled from 3 independent experiments). Unpaired t-test. p > 0.05.

Using this method, we assessed invasion, spread, and proliferation in the tissue after overlay. BRPKp110 invaded the LN tissue in the first hour such that they were not washed away during the wash step but were still rounded in morphology. By 20 hr, the cell morphology had changed to elongated, characteristic of cell adhesion and spread (Figure 3B), and they had penetrated to an average depth of 140 ± 17 μm into the LN tissue (Figure 3C). The cancer cells continued proliferating in the tissue, as staining for Ki-67 revealed a similar proportion of proliferating BRPKp110 cells after 20 hr in the LN tissue as in culture of BRPKp110 cells alone (Figures 3D, E; isotype controls shown in Figure S2A). To test the generalizability of this approach, we examined two additional cancer cell lines: HR+ B16F10 murine melanoma and HR- 4T1 murine mammary carcinoma cells. Both cell lines demonstrated the ability to infiltrate LN tissue, showing invasion after 1 hr and further spreading after 20 hr of culture (Figure S2B). Thus, live LN slices could support an ex vivo model of cancer cell invasion and spread across multiple cancer cell lines.

### Enrichment of cancer cells in the SCS preceded spread to the cortex and B cell follicle zones

Similar to direct intra-LN injection performed in vivo,^7,21^ adding cancer cells directly to the face of a LN slice allows the cells to bypass the afferent lymphatic vasculature. We took advantage of this feature to determine which regions of the LN were preferentially colonized by cancer cells in the absence of access barriers. To do so, we compared invasion between LN regions, using live tissue immunostaining and image segmentation to define the SCS, cortex, B cell follicles, T cell zone and medulla (Figure 4A). Invasion was normalized to the relative area of each zone to define an invasion-fold change, where a higher value indicated a greater cancer positive area per unit area of the region, and a value of 1 indicated a fractional cancer-positive area equal to the mean in the entire tissue slice.

**Figure 4.**
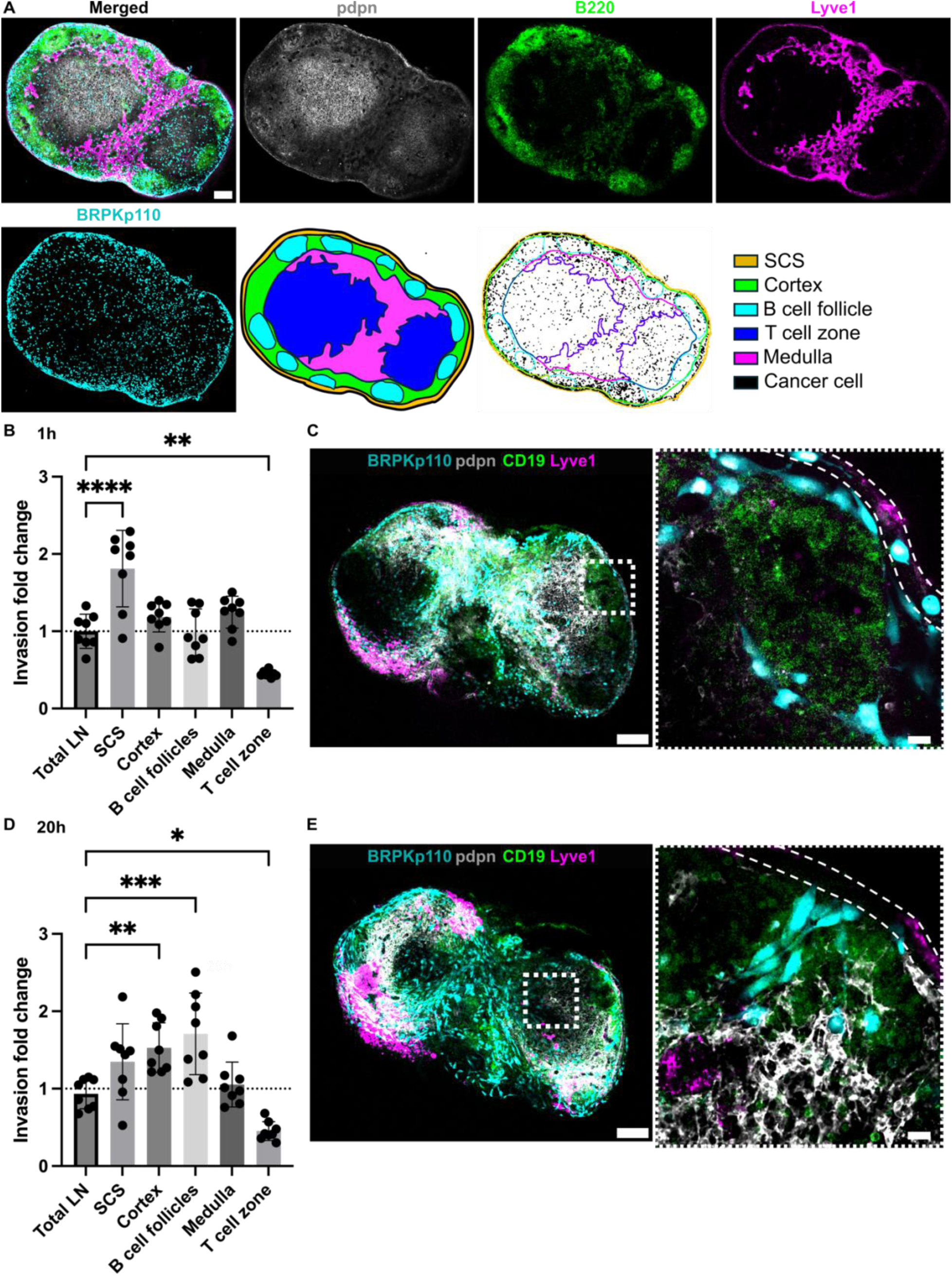
Dynamic distribution of cancer cells across LN zones. (A) Live immunofluorescence and image segmentation strategy for quantification of cancer cell invasion in LN zones. Representative images of LN tissue slice that was overlaid with cancer cells (NHS-Rhodamine, cyan) and stained for podoplanin (pdpn, gray), a B cell marker (B220, green), and lymphatic endothelial cells (lyve1, magenta). Result of image segmentation for assignment of LN regions. (B, D) Invasion fold change of BRPKp110 cells across LN zones 1 hr (B) and 20 hr (D) post-seeding, normalized to the average invasion of the total LN area. Mean ± stdev; each data point represents one LN slice (n = 7-8/per group, pooled from 3 mice). One-way ANOVA, followed by Dunnett posthoc test. *p &lt;0.05, **p &lt; 0.01, ***p = 0.001, ****p &lt; 0.0001. (C, E) Representative images of BRPKp110 cells (cyan) invasion in the SCS at 1h (C) and cortex and B cell follicles at 20 hr (E) post-seeding. Scale bars: left image 200 µm; right image 20 µm. LN tissues were stained with a B cell marker (CD19, green) and lymphatic cell marker (Lyve-1, magenta).

We assessed the distribution of the cancer cells at 1, 20, and 40 hr after seeding, hypothesizing that there would be reorganization over time. At 1 hr after seeding, there was a notably greater distribution of BRPKp110 cells within the SCS and significantly lower in T cell zone in comparison to the average across the tissue (Figure 4B). Indeed, individual cancer cells were clearly visible inside the SCS (Figure 4C), as well as elsewhere in the tissue. However, by 20 hr after seeding, the enrichment of BRPKp110 cells within the SCS was no longer statistically significant; instead, cancer cells were preferentially distributed within the cortex and B cell follicles. No difference was detected in the regional distribution of cancer cells between the 20 hr and 40 hr culture periods (Figure S3). Thus, cancer cells initially entered the tissue preferentially in the SCS, followed by a re-distribution into the cortex and B cell zones, with relative exclusion from the central T cell zones at both times. This behavior was reminiscent of the in vivo behavior of melanoma cancer cells in TDLN, where metastatic cells first accumulated in the SCS in response to a CCL1 gradient and later formed metastatic lesions in the deeper parenchyma.^13^

### Ex vivo invasion correlated with the distribution of CXCL13 and CCL1 in naïve LN slices

Chemokines establish both soluble and immobilized concentration gradients. To define which zones of naïve LNs expressed immobilized CCL21, CCL1, CXCL12 and CXCL13 and how these changed during LN slice culture, we used live immunofluorescence labeling (Figure 5A) and image segmentation as in Figure 2A. The distribution of immobilized CCL21 and CXCL13 in LN culture exhibited dynamic changes over time from 1 to 20 hr (Figure 5B), with a significant decrease in CCL21+ area (76% decrease, p < 0.001) and an increase in CXCL13+ area (94% increase, p < 0.01). No changes in the CCL1+ or CXCL12+ area was detected in this time. None of the chemokines were confined to a specific anatomical zone of LN, but rather were distributed across all anatomical zones of the LN to varying degrees (Figures S4A, B).

**Figure 5.**
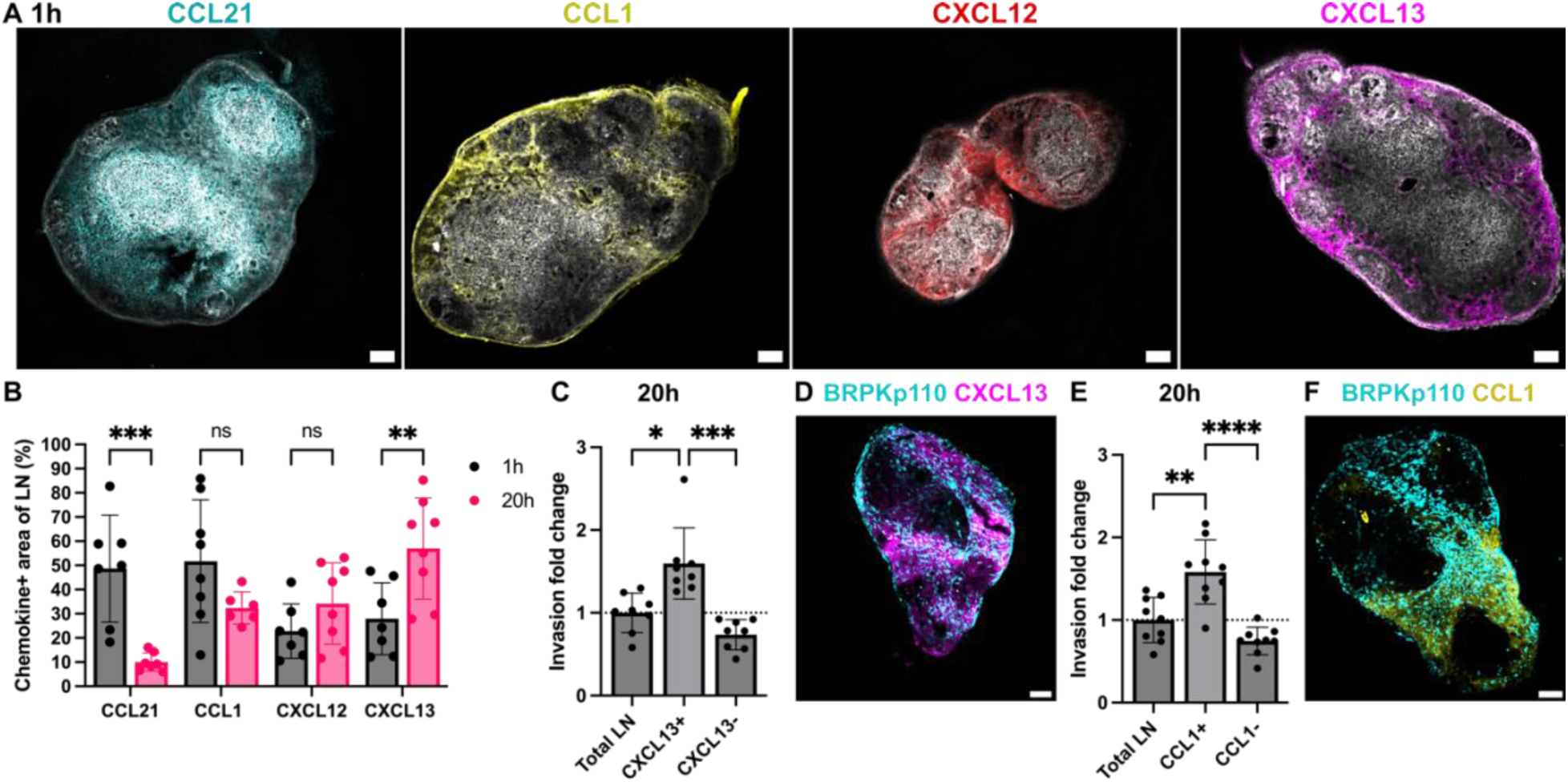
Spatiotemporal invasion of cancer cells in regions of immobilized chemokines. (A) Representative images of LN stained for podoplanin (pdpn, gray) and immobiloized chemokines: CCL21 (cyan), CCL1 (yellow), CXCL12 (red) and CXCL13 (magenta) chemokines after 1 hr of culture. (B) Fraction of LN area positive for CCL21, CCL1, CXCL12 and CXCL13 after 1 hr and 20 hr of culture. Mean ± stdev; each data point represents measurement from one LN slice (n = 7- 8/per group, LN slices obtained from 3 mice). Two-way ANOVA with Sidak posthoc test. *p &lt; 0.05, ***p &lt; 0.001. (C, E) Invasion fold change of BRPKp110 cells in chemokine positive and negative regions of the LN, normalized to the average invasion of the total LN area, after 20 hr of culture. CXCL13 in (C) and CCL1 in (E). Mean ± stdev; each data point represents one LN slice (n = 7-8/per group, LN slices obtained from 3 mice). One-way ANOVA with Tukey posthoc test. *p &lt; 0.05, ***p &lt; 0.001. (D, F) Representative images of BRPKp110 cells in LN slices with immunolabelling for CXCL13 (D) and CCL1 (F) after 20 hr of culture. Scale bars 200 µm.

As the chemokines were distributed throughout the LN, we next asked the extent to which BRPKp110 cancer cell invasion in this ex vivo model correlated with distribution of immobilized chemokines. Cancer cell invasion within chemokine-positive and chemokine-negative regions was compared to the average invasion across the LN slice. To avoid neutralizing any chemokine activity, immunofluorescence labeling was performed after cancer cell invasion in these experiments. At 1 hr post-seeding, BRPKp110 invasion was 1.6-fold higher in the CXCL13+ region compared to the tissue average (Figures S4B, C). After 20 hr of culture, invasion rate remained high in the CXCL13+ region (1.5-fold increase over the average) and was also increased in the CCL1+ region (1.3-fold increase over the average) (Figures 5C-F). No enrichment was detected in other chemokine-positive or negative regions at either time point (Figure S4D). Thus, we established a correlation between spatiotemporal invasion of BRPKp110 cancer cells in naïve LN tissue and distribution of immobilized CXCL13 and CCL1. Considering that the chemokines were detected across multiple zones of the LN, we concluded that cancer cell distribution was better predicted by the distribution of chemokine-rich domains than by anatomical zone.

### Knock-out of CXCR5 in BRPKp110 impaired migration into the LN and revealed redundancy in chemotactic migration

A feature of the ex vivo model is that it isolates the impact of changes in cancer cell signaling on invasion of the LN, without confounding effects from changes to migration out of the primary tumor or entry or migration through the lymphatic vasculature. Having found preferential BRPKp110 invasion towards CXCL13 at both 1 hr and 20 hr after overlay, we sought to demonstrate this capability by testing the requirement for the cognate chemokine receptor, CXCR5, in facilitating localization in the LN. We utilized CRISPR (clustered, regularly interspaced, short palindromic repeats)/Cas9 (CRISPR-associated protein 9) technology to generate BRPKp110 cell lines lacking function of CXCR5. To facilitate interaction with Cas9, we employed chemically modified synthetic CXCR5 gene–specific CRISPR RNAs (crRNA) along with fluorescently labeled tracer RNAs (tracrRNAs), enabling the selection of the transfected population through cell sorting (Figure 6A). After transfection, the viable fraction of tracrRNA- positive BRPKp110 cells, which constituted 85.5% of all cells, was isolated and cultured to establish the BRPKp110 CXCR5 knockout (KO) cell line (Figure 6B). We confirmed the loss of chemotactic function in CXCR5 KO cells using a 3D transwell assay with media supplemented with CXCL13 (Figure 6C). CCR7 KO cells were generated and validated as well (Figures S5A, B). CCR8 KO cells were also produced, but they retained chemotactic function towards CCL1 and were not pursued further.

**Figure 6.**
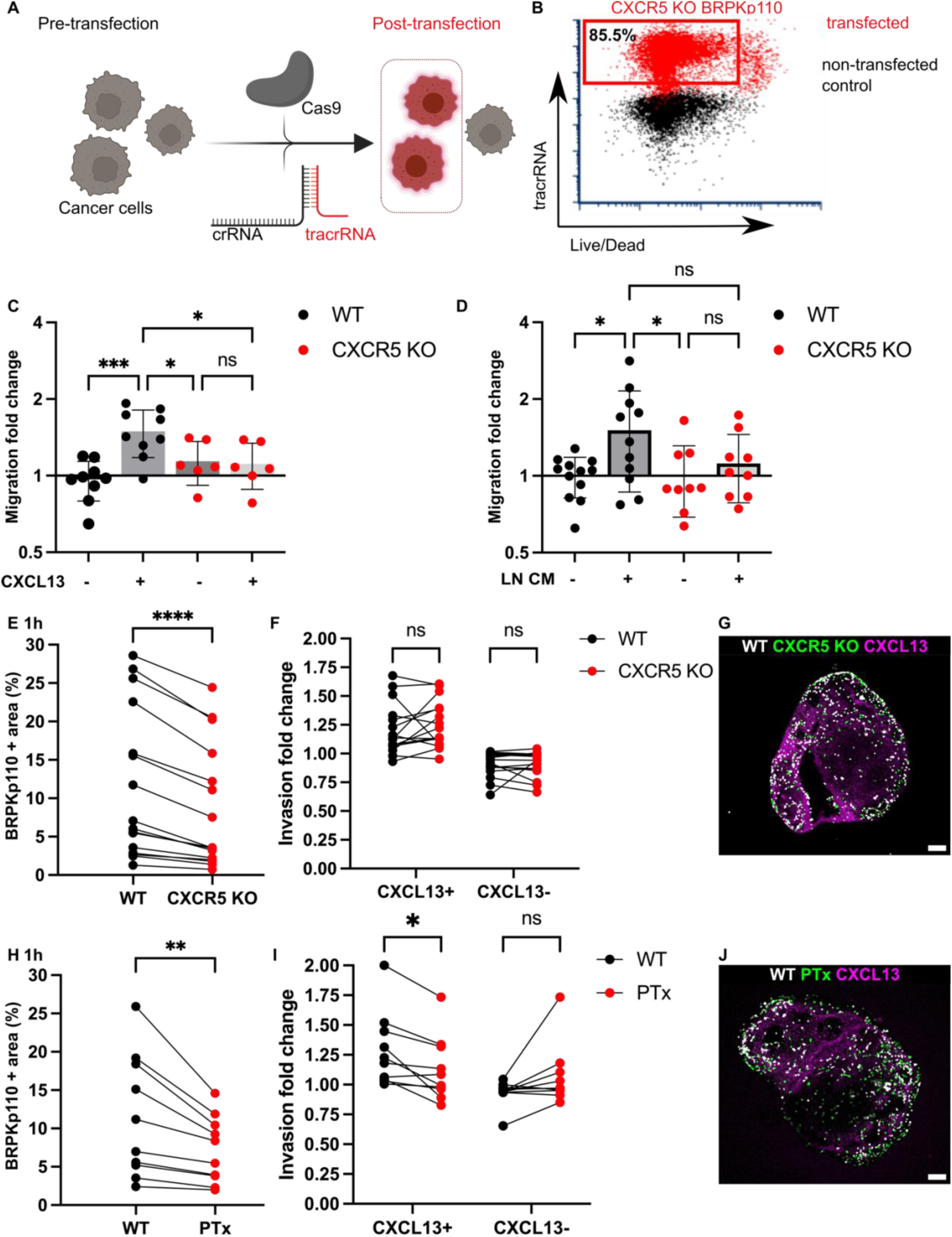
Blockade of CXCR5 mediated signaling alone was not sufficient to prevent cancer cell chemotactic migration into LN tissue. (A) Application of CRISPR/Cas9 technology for generation of cancer cell lines lacking CXCR5. crRNA, fluorescently labeled tracrRNA, and recombinant Cas9 protein. (B) TracrRNA signal used to select the transfected population in CRISPR-treated cells (red). Non-transfected control WT BRPKp110 (black) shown for comparison. (C) CXCR5 KO migration toward media containing 200 ng/mL of CXCL13 was impaired, confirming the loss of receptor function. Each data point represents the mean migration fold change per membrane, calculated from three non-overlapping fields of view (n = 2-3 membranes/condition; normalized data pooled from 3 independent in vitro experiments). Two-way ANOVA with Tukey posthoc test. ***p &lt; 0.0001, *p &lt; 0.05. (D) Migration fold change of WT and CXCR5 KO BRPKp110 cells towards media conditioned by culture of naïve LN CM from culture. Each data point represents the mean migration fold change per membrane, calculated from three non-overlapping fields of view (n = 3-4 membranes/condition; normalized data pooled from 3 independent in vitro experiments). Two-way ANOVA with Tukey posthoc test. *p &lt; 0.05. (E, H) Fraction of total LN area positive for cancer cells after 1 hr overlay. Paired comparison between WT and CXCR5 KO BRPKp110 (E) and WT and PTx treated BRPKp110 (H). Each data point represents paired measurements from one LN slice (n = 10-14/per group, LN slices obtained from 3 mice). Paired t-test. ***p &lt; 0.001, **p &lt; 0.01 (F, I) Invasion fold change of cancer cells in CXCL13+ domain after 1 hr post overlay. Paired comparison between WT and CXCR5 KO BRPKp110 (F) and WT and PTx treated BRPKp110 (I). Each data point represents invasion fold per LN slice (n = 10-14/per group, LN slices obtained from 3 mice). Two-way ANOVA, followed by Sidak posthoc test. *p &lt; 0.5. (G, J) Representative image of cancer cells distribution after 1h post seeding in naïve LN labeled for CXCL13 (magenta). (G)WT BRPKp110 (gray) and CXCR5 KO (green). (J) WT BRPKp110 (gray) and PTx-treated (green). Scale bars 200 µm.

First, we tested requirement for CXCR5 in cancer cell migration towards factors secreted by naïve LN in vitro, by using conditioned media obtained from overnight culture of naïve LN slices in a 3D transwell assay. The mean change in migration towards LN CM was 26% reduced in CXCR5 KO as compared to wild type (WT) BRPKp110 (Figure 6D). On the other hand, there was substantial within-group variation between supernatants from different slices, leaving the migration towards CM not significantly different between WT and KO cells. This result suggested that targeting the CXCR5 receptor reduced the migration of cancer cells toward factors secreted by naïve LN, but perhaps did not completely eliminate it.

Next, we tested the requirement of CXCR5 for cancer cell invasion into naïve LN tissue, and into the CXCL13+ domain in particular. To allow paired comparisons of invasion, we overlayed equal numbers of CXCR5 KO and WT BRPKp110 cells, labeled with different fluorophores, onto each LN slice. In line with the in vitro results, we found that CXCR5 KO cells invaded less into each slice than the WT cells (35% mean reduction in invasion; Figure 6E), though some cells did still enter the tissue. Interestingly, although total invasion was reduced, invasion of the CXCL13+ domain was unaffected by KO of CXCR5 alone (Figure 6F). Only complete blockade of chemokine signaling by PTx treatment significantly reduced the BRPKp110 invasion in the CXCL13+ regions (Figures 6G, H), an effect that remained after 20 hr of culture (Figure S5C). Thus, we concluded that the migration of CXCR5 KO cells towards CXCL13+ regions was driven by chemotaxis towards other chemokines.

These findings collectively suggested that CXCR5 was required for a portion of the total BRPKp110 invasion into naïve LNs, but that disrupting CXCR5-mediated signaling alone was insufficient to prevent invasion towards domains rich in CXCL13, due to the multiple chemokines expressed in any given region. These experiments were enabled by the isolation of the LN in the ex vivo model and would be challenging to conduct in vivo, since CXCL13/CXCR5 axis also plays a substantial role within the tumor itself.^49,50^

### Primary pre-metastatic TDLNs experienced reduced initial invasion of cancer cells despite increased chemokine secretion

Having established the model of cancer cell invasion in naïve LN slices, we proceeded to apply this model to predict invasion dynamics within the pre-metastatic TDLN in breast cancer.

Standard in vivo experiments are complicated by the fact that the tumor and TDLN co-evolve.

Therefore, here we applied the ex vivo model of invasion to address whether identical cancer cells invaded differently into pre-metastatic TDLN vs naïve LN.

To generate TDLN, we used a well-established murine model of breast cancer, in which BRPKp100 cells were inoculated into the fourth abdominal mammary fat pad on each side of the animal (Figure 7A). In this model, the inguinal TDLN (iTDLN) and axillary TDLN (aTDLN) represent the primary and secondary TDLNs, respectively.^51^ TDLNs were harvested at day 5 post tumor inoculation, a timepoint preceding palpable tumor formation (Figure 7B), when no BRPKp110 cells (anti-GFP+ CD45-) were detectable in the TDLNs via flow cytometry (Figure 7C, Figure S6A). Therefore, we considered this timepoint to be pre-metastatic, although we cannot exclude the presence of a small, undetectable number of cells or tumor-derived fragments.

**Figure 7.**
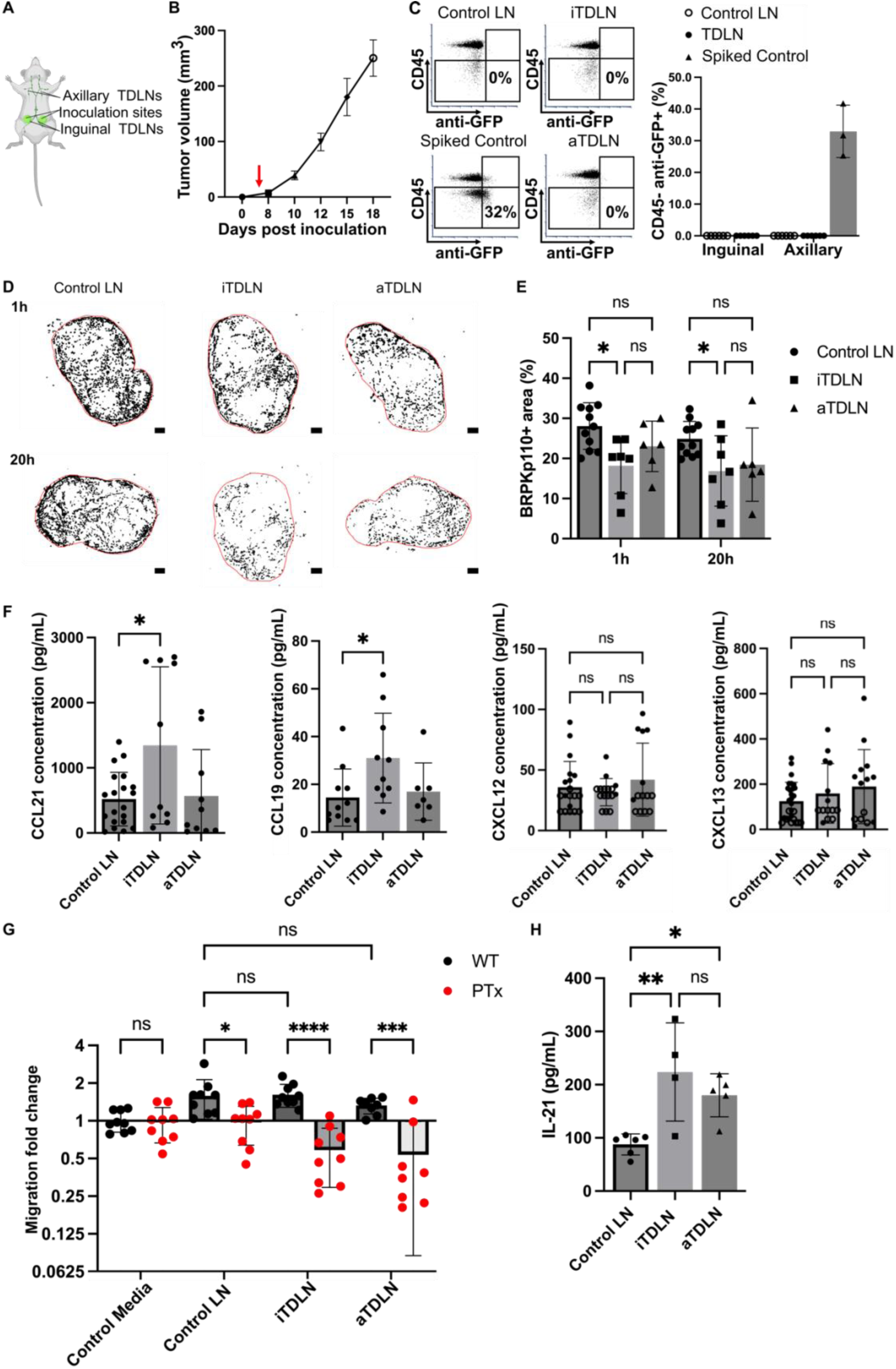
**Reduced invasion of cancer cells in pre-metastatic iTDLN ex vivo**. (A) Schematic illustration of in vivo model of breast cancer from which TDLN were obtained. Bottom-up view of the animal. (B) Growth kinetics of BRPKp110 mammary tumors (n = 3 mice). (C) Flow cytometry analysis of cancer cell in TDLN. Quantification of CD45- anti-GFP+ cells in TDLNs 5 days post BRPKp110 inoculation. Mean ± stdev; each data point represents a fraction of CD45- anti-GFP+ cells per LN (n = 6 LNs/ group (inguinal, axillary) obtained from 3 tumor-bearing mice and 3 control mice injected with PBS). Two-way ANOVA, followed by Tukey posthoc test. p > 0.05. (D) Representative images of cancer cell invasion (WT BRPKp110, black) into control LN, pre-metastatic iTDLN and aTDLN at 1 hr and 20 hr post overlay. Scale bar 200 µm. (E) BRPKp110+ area positive area in control LNs, a non-tumor mice injected with PBS, pre-metastatic iTDLN and aTDLN after 1h and 20h of culture. Mean ± stdev; each data point represents an individual LN slice (n=2-3/group, LN slices obtained from 3 mice). Two-way ANOVA with Tukey posthoc test. *p &lt; 0.05. (F) Concentrations of CCL21, CCL19, CXCL12 and CXCL13 in CM from TDLN slices after 20 hr culture, measured by ELISA. Mean ± stdev; each dot shows the supernatant from one LN slice (n = 8-10 slices, pooled from 3 female mice). Two-way ANOVA with Tukey posthoc test. *p &lt; 0.05. (G) Migration of untreated and PT-treated BRPKp110 towards TDLN CM. Mean ± stdev; each data point represents the mean migration fold change per membrane, calculated from three non-overlapping fields of view (n = 3-4 membranes/condition; normalized data pooled from 3 independent in vitro experiments). Two-way ANOVA, followed by Tukey posthoc test. *p &lt; 0.05, ***p &lt; 0.001, ****p &lt; 0.0001. (H) Intranodal levels of IL-21 in were significantly higher in pre-metastatic iTDLN and aTDLN than in control LN. Mean ± stdev; each data point represents the contents of 2 pooled LNs. Control: 12 LNs, 3 mice. iTDLN: 8 LNs, 4 mice, aTDLN: 10 LNs, 5 mice. Two-way ANOVA, followed by Tukey posthoc test. **p &lt; 0.01, *p &lt; 0.05.

To compare the invasion potential of pre-metastatic TDLN versus control LN, we seeded BRPKp110 cells from cell culture onto the day-5 ex vivo slices of TDLN or control LN from PBS-injected animals. As in naïve LN, cancer cells readily entered the TDLN slice ex vivo and converted from a round to spread morphology between 1 and 20 hr (Figure 7D). Strikingly, the fraction of LN area occupied by cancer cells was significantly lower in iTDLN slices compared to control LNs (Figures 7D, E). This reduction was observed both at the initial entry (25% decrease) and after 20 hr (19% decrease), suggesting less initial accumulation rather than reduced survival or proliferation in overnight culture.

To attempt to determine the origin of the reduced invasion into TDLN, we first tested whether levels of secreted chemokines were similarly reduced. However, overnight cultures of primary draining iTDLN tissue slices actually secreted significantly more CCL21 and CCL19 into the supernatant compared in comparison to aTDLN and control LN (Figure 7F), with a correlation between CCL19 and CCL21 secretion only in the iTDLNs (Figure S6B). The secretion of CXCL12 and CXCL13 by TDLN was not different from that of LNs obtained from control mice (Figure 7F), and immunofluorescence labeling revealed no differences in the fractions of area positive for immobilized chemokines between TDLNs and control LNs (Figure S6C). Thus, the reduced invasion of cancer cells into iTDLN slices was not attributable to reduced secretion of secreted or immobilized chemokines, as secretion was unchanged or even increased. In agreement with these data, BRPKp110 cells showed similar migration in transwell assays towards media conditioned by pre-metastatic TDLNs as by naïve LN (Figure 7G). The migration was abolished by PTx treatment (Figure 7G) and was reduced in CXCR5 and CCR7 KO cells similarly to in WT cells (Figure S6D). Together these data confirmed that chemotaxis was intact towards TDLN conditioned media.

Next, we considered that reduced invasion might result from anti-tumor immunity in the pre-metastatic TDLN. Recent studies have highlighted the emerging role of interleukin-21 (IL-21) in the immune response against breast cancer in humans and mouse models. In breast cancer patients, elevated levels of IL-21 in CD4+ T cells were linked to better prognostic outcomes.^52^ Additionally, in a murine model of 4T1 breast cancer, elevated IL-21 was identified as a crucial regulator of CD8+ T-cell-mediated antitumor immunity in the pre-metastatic TDLN.^21^ In line with those reports, we observed significantly increased levels of intranodal IL-21 in pre-metastatic (day 5) iTDLNs and aTDLNs from the BRPKp110 animals compared to PBS control animals (Figure 7H). Thus, reduced invasion of cancer cell into iTDLN correlated with increased intranodal levels of IL-21, consistent with potential immune activation. We plan to explore the immune state of the TDLN further in future work.

In summary, the ex vivo LN slice model predicted a lower invasion potential of pre-metastatic iTDLNs compared to control LNs, which was not due to diminished chemokine secretion, and which correlated with elevated intranodal IL-21 in concordance with prior reports. Understanding the mechanism behind reduced invasion remains a key focus for future research.

## DISCUSSION

This work showed that live LN tissue slices form a powerful system to model cancer cell spread within the complex LN microenvironment ex vivo, in the absence of lymphatic barriers. LN slices secreted multiple chemokines that attracted cancer cells into the tissue both individually and in the conditioned media. This LN slice model was well suited to quantify the capacity of cancer cells to invade and colonize distinct regions of the LN, and it predicted a dynamic invasion process. Accumulation occurred initially preferentially in the SCS, followed by subsequent spread to the cortex and B-cell follicles, similar to published in vivo reports.^53^ The distribution of cancer cells within the LN correlated with the distribution of immobilized CXCL13 and CCL1 chemokine-rich domains within the LN. Blocking an individual chemokine receptor led to reduced overall invasion but was not sufficient to diminish cancer cell enrichment; this result confirmed that the ex vivo model was capable of supporting complex and overlapping signaling pathways. Furthermore, this system was readily applied to TDLNs, where it predicted a lower invasion potential of cancer cells into pre-metastatic iTDLNs than naïve nodes, in line with other models of breast cancer.^21^

Overall, these results indicate that this novel ex vivo model is suitable for mechanistic analysis of tumor invasion and translational studies for drug testing. The model enhances experimental accessibility compared to in vivo models of TDLN metastasis, allowing simultaneous observation of the histological appearances and verification of the biological hypotheses. Furthermore, the model allows for the manipulation of cancer cells and LN slices in isolation, enabling the testing of cancer cell invasiveness into the LNs at various stages of disease progression in a controlled setting. Compared to existing 3D cell systems, the intact cellular organization and chemotactic function of live LN slices allow for the simultaneous assessment of molecular factors secreted at the tissue level and the effects of complex microenvironmental cues on cancer cell invasion patterns.

Several areas remain to improve the ex vivo model of LN metastasis in the future, and to use it to address additional questions. While we demonstrated that live LN slices effectively support the invasion of cancer cells for up to 20-48 hr of culture, this time period does not fully capture the long-term interactions and progressive stages of cancer cell invasion and colonization of the TDLN that occur in vivo. Future studies will aim to extend the culture duration. Furthermore, in contrast to in vivo models, where primary tumor progression is influenced by changes in the tumor microenvironment,^54^ our study used a cancer cell line under stable culture conditions. However, similar to the in vivo scenario where only a few cancer cells show metastatic potential, in our ex vivo set up we also observed that only a fraction of seeded cells invaded the LN tissue. To our knowledge this model is the first to enable study of cancer cell invasion in the complex cellular microenvironment of LN tissue, including myeloid cell populations that had relocated to TDLNs prior to collection of the LN tissue. However, this system does not replicate the new recruitment of migratory myeloid cell population into TDLN during culture. This aspect may be interesting for studying cancer cell invasion over extended culture periods. Future studies are needed to address these limitations.

The absence of lymphatic barriers in the live LN slice is both a strength and a limitation. Unlike in vivo models, delivery of cancer cells directly to the open face of the slice means that extravasation into the lymphatic vessels and out through the LN sinus floor is not required for entry into the LN. Those events are successfully mimicked by other models.^24,55–59^ In contrast, this ex vivo model specifically focuses on the events of cancer cell colonization of the LN parenchyma, to learn where and how cells accumulate and spread once barriers are disrupted. In this way it is a direct parallel to recent work that used in vivo injection of cancer cells into blood vessels to identify favorable niches for metastasis.^60^ Interestingly, despite the different entry route, cancer cells in this model still favored initial invasion of the LN SCS, similar to in vivo results.^20,53,61^

Looking forward, we anticipate that this ex vivo model of LN metastasis will enable a host of future studies. In addition to mechanistic studies of cancer cell invasion into LNs of varied inflammatory and cancer-primed states, the model is also potentially suitable to study TDLN- induced cancer cell damage or death. Furthermore, with their intact immune function,^33^ LN slices may serve as an excellent model to test the impacts of TDLN-induced immunosuppression, as hinted at in prior work in on-chip co-cultures.^26^ The ability to mix and match cancer cells and LN tissue from various stages of cancer progress, as well as from different drug treatments, ages, and comorbidities, makes the model uniquely complementary to in vivo studies, with many potential applications.

## MATERIALS AND METHODS

### Cell culture

Mouse mammary cancer cell lines BRPKp110-GFP+, 4T1-luc-red and melanoma B16F10 were obtained from Melanie Rutkowski, University of Virginia. Cells were cultured in RMPI (Gibco, 2505339) supplemented with 10% FBS (Corning Heat-inactivated, USDA approved origin, lot: 301210001), 1x L-glutamine (Gibco Life Technologies, lot: 2472354), 50 U/mL Pen/Strep (Gibco, lot: 2441845), 50 μM beta-mercaptoethanol (Gibco, 21985-023), 1 mM sodium pyruvate (Hyclone, 2492879), 1× non-essential amino acids (GIBCO, 2028868), and 20 mM HEPES (Gibco, 15630-060). Cells were seeded in T75 or T175 flasks (Nunc™ EasYFlask™, Fisher Scientific) following manufacturer’s recommendations on seeding cell density and cultured sterilely in humidified atmosphere of 5% CO2 and 95% oxygen at 37°C. The cells were passaged upon reaching 70–80% confluence with 0.25% trypsin/EDTA (Invitrogen, ThermoFisher Scientific) with a 1:4 split ratio. All cell lines were maintained for less than four passages, with monitoring of morphology and testing for mycoplasma.

### Animal work

All animal work was approved by the Institutional Animal Care and Use Committee at the University of Virginia under protocol no. 4042 and was conducted in compliance with guidelines the Office of Laboratory Animal Welfare at the National Institutes of Health (United States). C57BL/6 mice ages 6–12 weeks (Jackson Laboratory, U.S.A.) were housed in a vivarium and given water and food ad libitum. Due to the prevalence of the breast cancer in women, only female mice were used in this study. For generation of tumors in vivo, 5· 𝟏𝟎^𝟓^ BRPKp110 cells were suspended in 100 µL PBS and injected orthotopically into the abdominal mammary fat pad. A control group of female C57Bl/6 mice of matched age received an injection of PBS. Tumor size was measured by calipers every 2–3 days after reaching a palpable size.

### Generation of lymph node tissue slices

Lymph nodes were collected and sliced according to a previously established protocol.^62^ Briefly, on the day of the experiment, animals were anesthetized with isoflurane followed by cervical dislocation. Inguinal and axillary lymph nodes were collected and placed in ice-cold PBS supplemented with 2% heat inactivated FBS. Subsequently, the lymph nodes were embedded in 6% low melting point agarose at 50°C and allowed to solidify. Agarose blocks containing the lymph nodes were obtained using a 10 mm tissue punch. Slices with a thickness of 300 μm were obtained using a Leica VT1000S vibratome. Following sectioning, the slices were promptly transferred to complete RPMI medium and incubated for a minimum of 1 hr before use.

### ELISA for analysis of cytokines and chemokines

Lymph node slices were cultured in complete RPMI media for 20 hr. Culture supernatant was collected and analyzed by sandwich ELISA assay using DuoSet ELISA development kit (R&D Systems, Inc., Minneapolis, MN, USA). ELISAs were for CCL21 (catalog no. DY457), CCL19 (DY440), CCL1 (DY845), CXCL12 (DY460) and CXCL13 (DY470) according to the manufacturer’s protocol. For measurement of intranodal IL-21 levels, inguinal and axillary lymph nodes were collected and carefully disrupted in 150 μL of ice-cold phosphate buffer, minimizing cell rupture.^63^ The suspension was centrifuged at 1,500 rpm for 5 min, and the supernatant was collected. Samples were analyzed by sandwich ELISA assay using DuoSet ELISA development kit for Il-21 (catalog no. DY594; R&D Systems, Inc., Minneapolis, MN, USA). In all cases, plates were developed using TMB substrate (Fisher Scientific), stopped with 1 M sulfuric acid (Fisher Scientific), and absorbance values were read at 450 nm on a plate reader (CLARIOstar; BMG LabTech, Cary, NC). To determine concentration of sample solutions, calibration curves were fit in GraphPad Prism 9 with a sigmoidal 4 parameter curve. Limit of detection (LOD) was calculated from the average of the blank plus 3× stdev of the blank.

### In vitro 3D transwell migration assay

In vitro migration assays were performed based on a protocol previously published by the Munson laboratory.^64^ 1·10^5^BRPKp110 cells were resuspended in a 100 µL hydrogel containing 2.0 mg/ml collagen type I (rat tail, Ibidi) and 1 mg/ml fibrinogen (BD Biosciences), then seeded into 12 mm diameter culture inserts with 8 μm pores (Millipore, Bellerica, MA). After gelation, 700 µL of chemoattractant or control media was added to the bottom compartment. To avoid generating fluid flow, the media outside of the insert was leveled with the medium inside by adding 100 µL of media on top of the gel. Cells were allowed to migrate during incubation in a humidified atmosphere of 5% CO2 and 95% oxygen at 37°C for 20 hr. After incubation, the gels in the upper chamber were removed with a cotton-tip applicator. The tissue culture inserts were fixed with 4% paraformaldehyde for 20 minutes at room temperature, washed with ice-cold PBS, stained with 300 nM DAPI for 30 minutes at room temperature, washed again with ice-cold PBS, and visualized by fluorescence microscopy. DAPI+ cells at the membrane surface were counted in three non-overlapping fields per well. Three technical replicates were averaged for each experimental run to yield a single biological replicate for statistical analysis. Cancer cell migration fold was calculated as previously described.^64^

### Ex vivo overlay of cancer cells onto live lymph node slices

After collection, lymph node slices were left to rest for at least one hour. 1·10^6^ BRPKp110 cancer cells were first stained with NHS-Rhodamine (Fisher Scientific) or Cell Trace (Fisher Scientific) for 20 minutes in a humidified sterile incubator at 37 °C with 5% CO2. Following the incubation period, excess fluorescent dye was removed by centrifugation. The cells were then resuspended in 1mL of complete culture media and incubated at 37 °C with 5% CO2 for 10 minutes to allow fluorescent reagent to undergo acetate hydrolysis. Lymph node slices were placed onto parafilm and covered with an A2 stainless steel flat washer (10 mm outer diameter, 5.3 mm inner; Grainger, USA), creating a 1 mm deep well over each lymph node tissue sample. For an overlay, a 20 μL of cancer cell suspension (2·10^4^ cells) was added into a washer on top of each LN slices and incubated for an hour at 37 °C with 5% CO2. Following the incubation period, excess cancer cells was rinsed with pre-warmed complete media for 30 minutes at 37 °C, changing the media every 10 minutes.

### Immunostaining of live lymph node slices

Upon collection, the slices were allowed to rest for one hour before being labelled for live immunofluorescence following a previously established protocol.^65^ Briefly, slices were Fc-blocked with an anti-mouse CD16/32 antibody (BioLegend, San Diego, CA) at a concentration of 25 μg/mL in 1x PBS with 2% heat-inactivated FBS (Gibco, Fisher Scientific) and incubated for 30 minutes in a humidified sterile incubator at 37 °C with 5% CO2. To stain, a 10 μL of antibody cocktail, containing antibodies at a concentration of 20 μg/mL, was added and the slices were incubated for an additional hour. Antibodies are listed in Table S1. Following staining, slices were washed with PBS for 30 minutes at 37 °C, refreshing the PBS every 10-15 minutes.

### Cas9/RNP nucleofection

The following protocol was adapted from a method published previously.^66^

#### crRNA selection

Three crRNAs were selected per target using the Benchling (www.benchling.com) online platform. The target area was limited to the first ∼40% of the coding sequence, and preference was given to guides targeting different regions within this area. On-target and off-target scores were evaluated using IDT and Synthego. Guides with the highest on-target and off-target scores were selected. crRNAs were ordered from Integrated DNA Technologies (www.idtdna.com/CRISPR-Cas9) in their proprietary Alt-R format (Table S2).

#### Preparation of crRNA–tracrRNA duplex

To prepare the duplex, each Alt-R crRNA and Alt-R tracrRNA (catalog no. 1072534; IDT) or Alt- tracrRNA-ATTO550 (catalog no. 1075928; IDTd) was reconstituted to 100 µM with Nuclease- Free Duplex Buffer (IDT). Oligos were mixed at equimolar concentrations in a sterile PCR tube (e.g., 10 µl Alt-R crRNA and 10 µl Alt-R tracrRNA). Oligos were annealed by heating at 95°C for 5 minutes in PCR thermocycler and the mix was slowly cooled to room temperature.

#### Precomplexing of Cas9/RNP

In a PCR strip, three crRNA–tracrRNA duplexes (3 µl equal to 150 pmol each, total of 9 µl) and 6 µl (180 pmol) TrueCut Cas9 Protein v2 (catalog no. A36499; Thermo Fisher Scientific) were gently mix by pipetting up and down and incubated at room temperature for at least 10 minutes.

#### Nucleofection

3·10^6^ BRPKp110 cells were resuspended in 20 µl primary cell nucleofection solution (P4 Primary Cell 4D-Nucleofector X kit S (32 RCT, V4XP-4032; Lonza). Cells were mixed and incubated with 15 µl RNP at room temperature for 2 minutes. The cell/RNP mix was transferred to Nucleofection cuvette strips (4D-Nucleofector X kit S; Lonza). Cells were electroporated using a 4D nucleofector (4D-Nucleofector Core Unit: Lonza, AAF-1002B; 4D-Nucleofector X Unit: AAF-1002X; Lonza), and EN-138 pulse code. After nucleofection, transfected cells were resuspended in prewarmed complete RPMI media and cultured overnight. The next day, tracrRNA+ cells were sorted on a BD Influx™ cell sorter using BD FACS™ Sortware software. After sorting cells were cultured for 3-5 days.

### Flow cytometry

Tumor-draining and control lymph nodes were homogenized using glass slides. Cancer cell dissemination in TDLNs was quantified using flow cytometry acquisition on a Guava easyCyte™ 8HT (Merck Millipore, Billerica, MA, USA). Cell suspensions were first stained with viability dye 7-AAD (AAT Bioquest, Sunnyvale, CA, USA), followed by blocking Fc receptors with anti- CD16/32 (clone 93, purified), and surface staining with anti-mouse CD45 (30-F11, PE). Cells were then permeabilized using buffer set (Invitrogen) and stained intracellularly with anti-GFP (FM264G, APC).

### Image acquisition

Transwell membranes were imaged on an AxioObserver 7 inverted fluorescence microscope with a 5X Plan-Neofluar objective. (Zeiss Microscopy, Germany).

All imaging of LN tissues slices was performed on a Nikon A1Rsi confocal upright microscope, using 400, 487, 561, and 638 lasers with 450/50, 525/50, 600/50, and 685/70 GaAsp detectors. Images were collected with a 4x/ 0.20 and a 40x/ 0.45 NA Plan Apo NIR WD objective.

### Image analysis

Images were analyzed in ImageJ (version 2.14.0/1.54g).^67^ First, autofluorescent noise from the individual image channels was subtracted, defined as the mean fluorescent intensity ± 1 stdev of respective fluorescent minus one (FMO) controls (n = 3 FMO control per experiment). After noise subtraction, regions of interest (ROI) were selected using the wand tracing tool and/or manually adjusted to reflect anatomical regions. The SCS ROI was defined as the area between podoplanin-positive LECs lining the ceiling and lyve1-positive LECs lining the floor of the SCS. The B-cell ROI was identified as the B220 or CD19 positive area; the B cell follicle ROI was identified as B220 or CD19 positive circular area within the cortex regions. The medullary ROI was defined as a lyve1 positive area in the paracortex of the LN. The T cell ROI was identified as the area of the LN excluding the SCS, cortex, B cell follicles, and medulla ROIs. All regions were non-overlapping, except for B cell follicle ROIs overlapping with the cortex region. Chemokine-rich domains were identified as CCL1, CCL21, CXCL12 or CXCL13 positive ROI, after defining a threshold. Cancer cell fluorescent signals were converted to binary, and the cancer cell positive area within the total LN and each LN region was measured. Cancer cell invasion was quantified as the cancer cell positive area of the total LN area. Invasion of the individual ROI was normalized to the relative area of each ROI to define an invasion-fold change (Equation 1), where a higher value indicated a greater cancer positive area per unit area of the ROI, and a value of 1 indicated a fractional cancer-positive area equal to the mean in the total LN area.

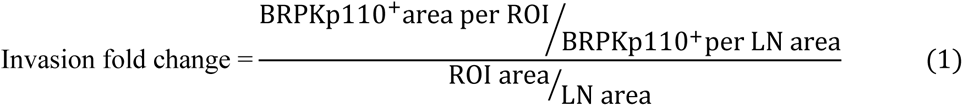

For representative image display, brightness and contrast were adjusted uniformly across all compared images within a figure unless otherwise specified.

### Statistical analysis

All in vitro assays were performed with a minimum of three biological replicates unless otherwise noted. Murine study numbers are noted in legends and by individual graphed data points. Graphs were generated using Graphpad Prism (version 9.4.0) software and were plotted with mean +/- stdev. p < 0.05 was considered statistically significant.

### Figure generation

Figures were generated using Inkscape (version 1.1). Schematics were generated using BioRender with license to RRP.

## DATA AVAILABILITY STATEMENT

Representative source data generated in this study are posted under Morgaenko et al. “**Ex vivo model of breast cancer cell invasion in live lymph node tissue**,” at https://dataverse.lib.virginia.edu/dataverse/PompanoLab.

## ETHICS STATEMENT

The animal study was reviewed and approved by University of Virginia Animal Care and Use Committee.

## AUTHOR CONTRIBUTIONS

Conceptualization: KM and RRP. Investigation: KM, AA, AGB, AMP, JMM, MRR, RRP. Formal analysis: KM. Data curation: KM. Project administration: KM and RRP. Resources: MRR and RRP. Software: RRP. Validation: KM, AA, AMP, JMM, MRR, RRP. Visualization: KM. Writing of original manuscript: KM and RRP. Review and editing of manuscript: KM, AA, AGB, AP, JMM, MRR, RRP. All authors contributed to the article and approved the submitted version.

## ACKNOWLEDGEMENTS

KM was supported in part by UVA Farrow Fellowship funding from the UVA Comprehensive Cancer Center. Research reported in this publication was additionally supported by the National Institute of Allergy and Infectious Diseases of the National Institute of Health under Award Numbers R01AI174207 and R01AI131723. The content is solely the responsibility of the authors and does not necessarily represent the official views of the National Institutes of Health. The authors thank Jennifer Hammel of Virginia Tech for guidance on the in vitro migration assay in culture inserts, and Morgan McKnight for technical assistance with LN tissue slicing.

