## Supporting information for "Ex vivo model of breast cancer cell invasion in live lymph node tissue"

### Contents

- Supplemental Figures S1-S6
- Supplemental Tables S1-S2

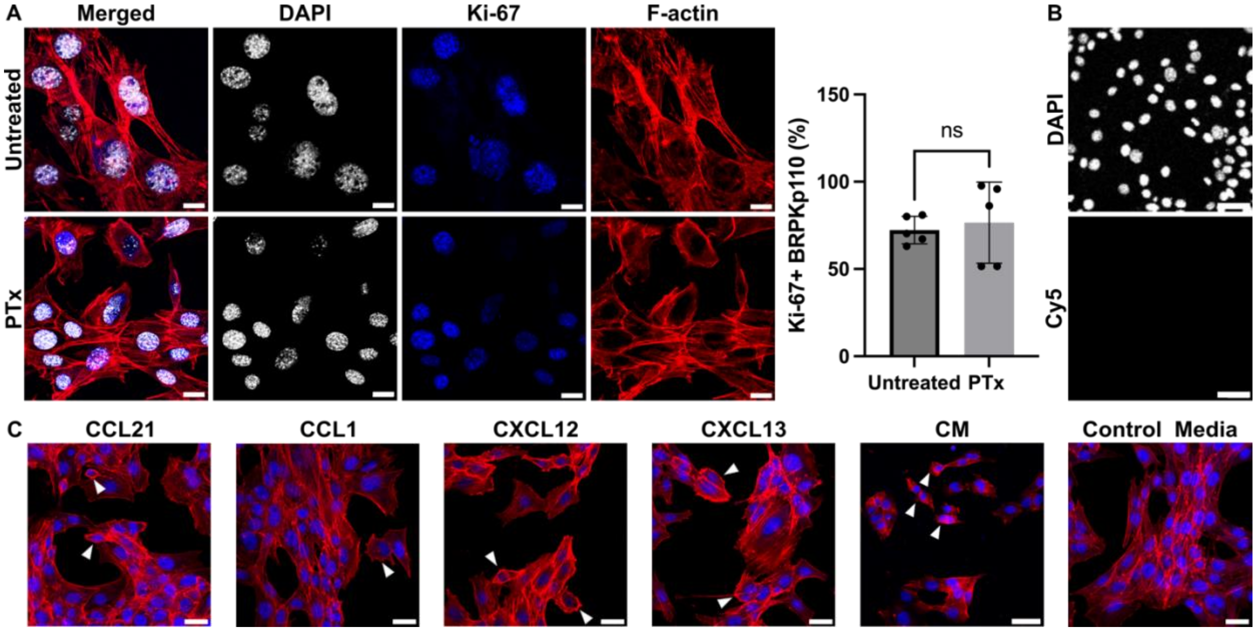


**Figure S1: Impact of PTx treatment and chemokines on BRPKp110 cells in culture.** (A) Morphology and proliferation of BRPKp110 cells cultured with 100 ng/ml PTx for 20 hr. Cells were stained with rhodamine-labeled phalloidin (red), Dapi (nuclei; white) and Ki-67 (blue). Graph shows the percent of Ki-67 positive cells per field of view in BRPKp110 cultured for 20h in media supplied with PTx or control media ("untreated"). Mean ± stdev; each data point represents a measurement from an individual sample (n=2-3/group, data pulled from 3 independent experiments). Unpaired t-test. $p > 0.05$. (B) Representative image of BRPKp110 cells stained with DAPI (nuclei; white) only, without receptor staining. Brightness and contrast are adjusted as in Figure 2F. Images were collected using DAPI and Cy5 channels. Scale bars 100 µm. (C) Effects of chemokines and conditioned media obtained from naïve LN culture on BRPKp110 cell morphology. Cells were stained with rhodamine-labeled phalloidin (red) and DAPI (nuclei; blue). Scale bars 20 µm.

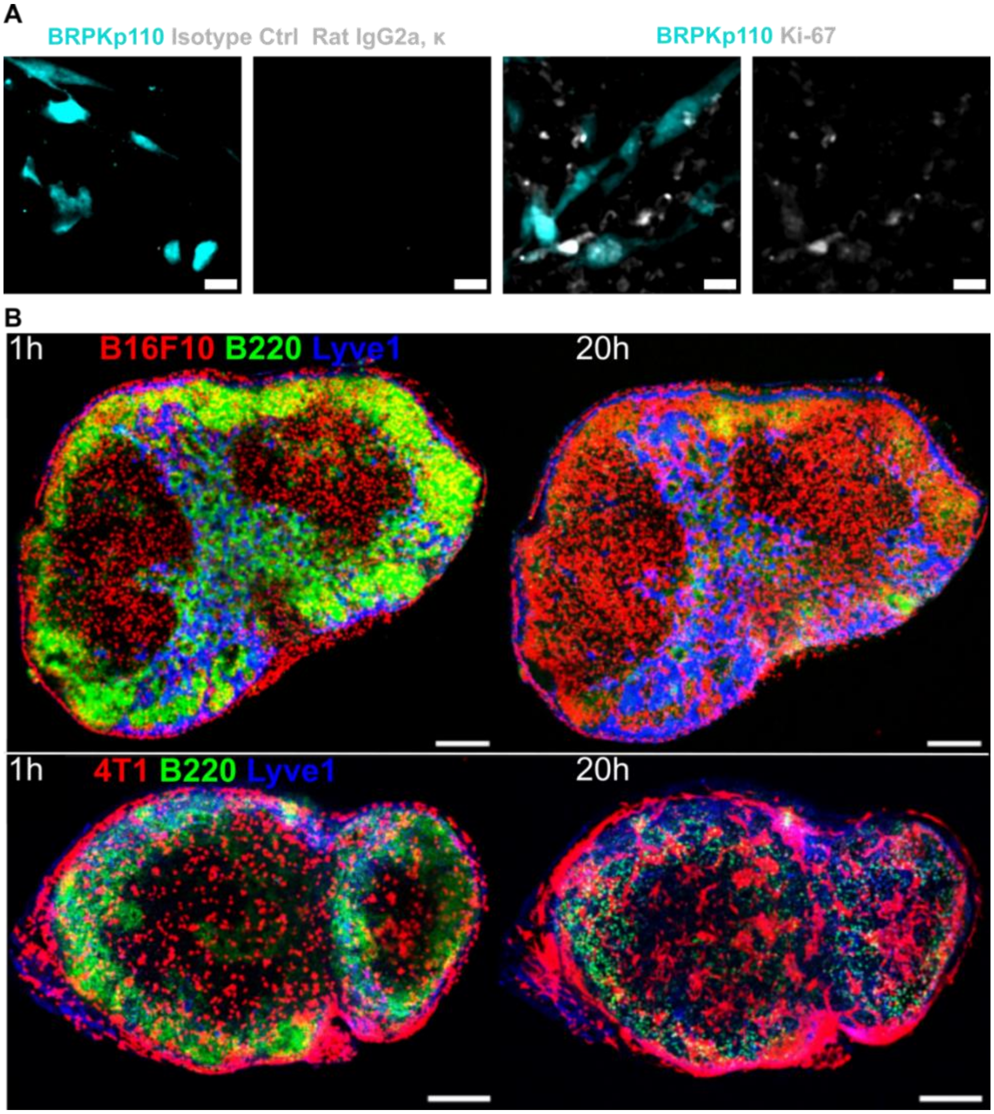


**Figure S2: Staining controls and infiltration patterns of cancer cells in ex vivo LN slices**. (A) Staining control experiment: BRPKp110 cells (NHS-Rhodamine, cyan) overlayed onto naïve LN slices after 20 hr of culture. *Left*: stained with an isotype control for the Ki-67 antibody, Rat IgG2a, k (gray). *Right*: stained with the Ki-67 antibody (gray). Scale bar 20 µm. (B) Infiltration of multiple types of cancer cells into ex vivo LN slices. Fluorescently labeled cancer cells (red) spread in naïve LN slices from 1 hour post seeding (left) to 20 hours of culture post seeding (right). *Top row*: B16F10 melanoma cells (red). *Bottom row*: 4T1 breast cancer cells (red). B220 (green) marks B cells, and Lyve1 (blue) marks LECs. Scale bar 200 µm.

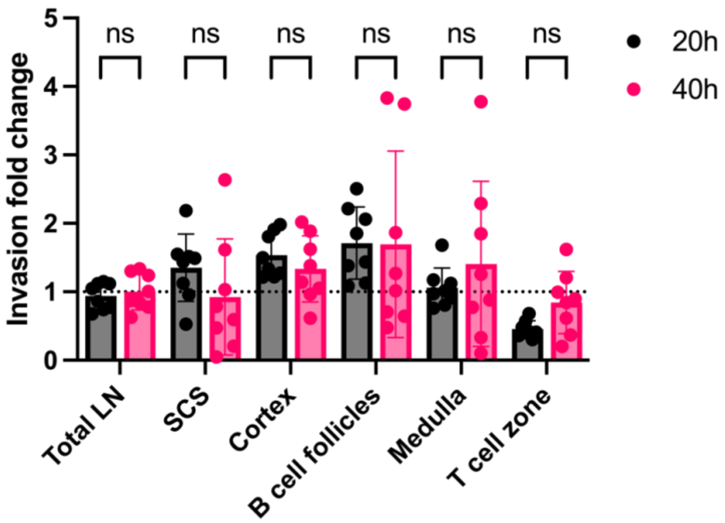


**Figure S3: Invasion dynamics of BRPKp110 cells in LN regions.** Invasion fold change of BRPKp110 cells in LN regions compared to the average across the LN tissue between 20 and 40 hrof culture. Data are presented as mean ± stdev. The dotted line represents normalized enrichment across the total area of the LN slice. Each data point represents the invasion fold change normalized to the total LN invasion on a per slice basis (n = 7-8 per group, LN slices obtained from 3 mice). Statistical analysis was performed using one-way ANOVA followed by Dunnett's posthoc test. $p > 0.05$.

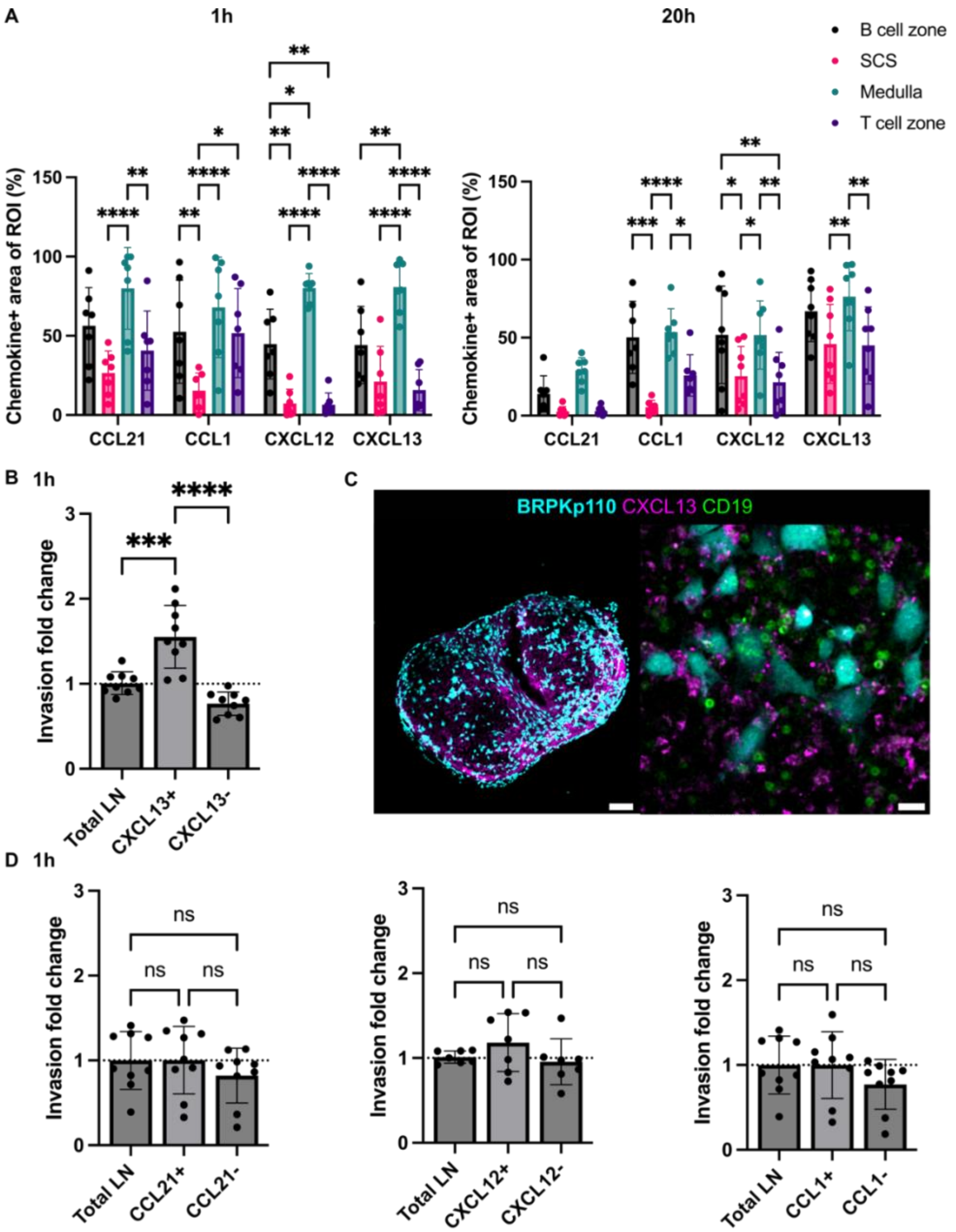


**Figure S4: Distribution of immobilized chemokine in regions of LN and BRPKp110 cell invasion in chemokine-rich domains.** (A)Fraction of total LN ROI positive for immobilized CCL21, CCL1, CXCL12 and CXCL13 measured in naive LN after 1 hr (left) and 20h (right) of culture. Mean ± stdev; each data point represents a LN slice (n=6-8/per group, LN slices obtained from 3 mice). Two-way ANOVA with Tukey posthoc test. *p < 0.05, **p < 0.01, ***p < 0.001, ****p < 0.0001.(B) BRPKp110 invasion fold in CXCL13 positive (CXCL13+) and CXCL13 negative (CXCL13-) regions of LN tissue relative to the invasion to the total LN tissue after 1 hr of culture. Mean ± stdev; dotted line represents normalized enrichment across the total area of the LN slice; each data point represents invasion fold change normalized to the total LN invasion on a per slice basis. One-way ANOVA, followed by Tukey posthoc test *p < 0.05, ***p < 0.001. (C) Representative images of cancer cells (cyan) in CXCL13+ chemokine domain (magenta) after 1 hr of culture. Scale bars 200 µm (left), 20 µm (right). (D) BRPKp110 invasion fold in chemokine-rich domains of LN tissue relative to the invasion to the total LN tissue after 1 hr of culture. Mean ± stdev; each data point represents a LN slice (n=7-9/per group, LN slices obtained from 3 mice). One-way ANOVA, followed by Tukey posthoc test. p > 0.05.

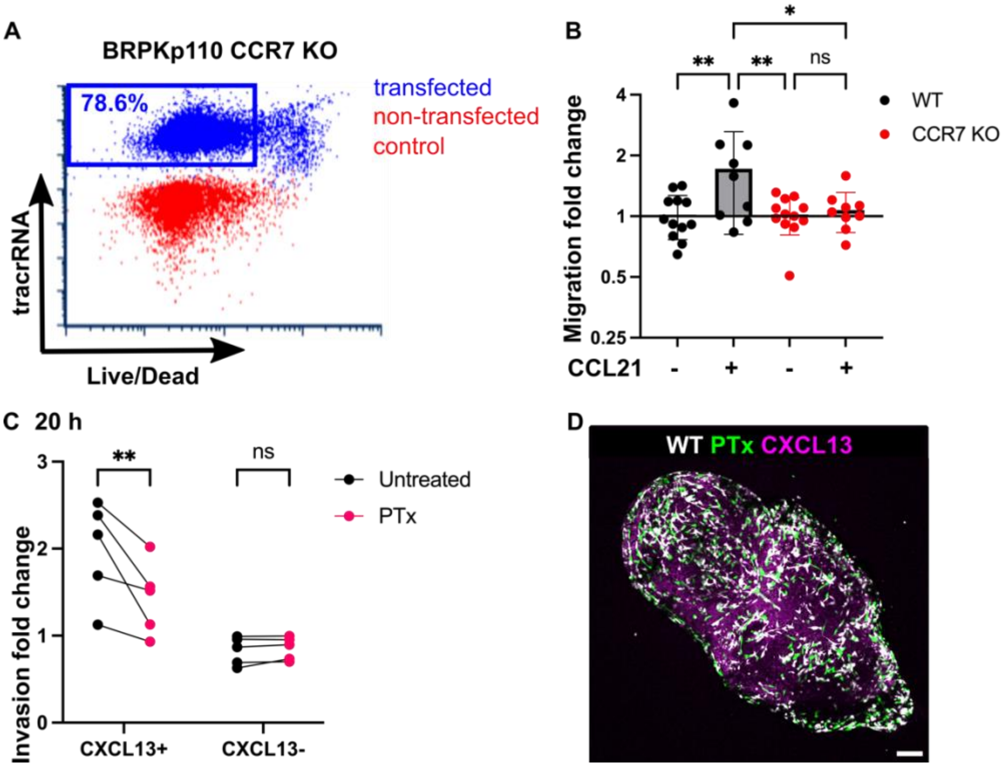


**Figure S5: Impaired chemotactic function in BRPKp110 CCR7 knockout and effects of PTx treatment on BRPKp110 invasion in LN slices.** (A) Selection of tracrRNA positive cell population post-transfection (blue), non-transfected control WT BRPKp110 (red). (B) CCR7 KO migration toward media containing 200 ng/mL of CCL21 was impaired, confirming the loss of receptor function. Mean ± stdev; each data point represents the mean migration fold change per membrane, calculated from three non-overlapping fields of view (n = 2-3 membranes/condition; normalized data pooled from 3 independent in vitro experiments). Two-way ANOVA, followed by Tukey posthoc test. **p < 0.01, *p < 0.05. (C) Invasion fold change of untreated WT BRPKp110 and PTx pre-treated BRPKp110 in CXCL13+ domain after 20 hr post overlay. Mean ± stdev; each data point represents invasion fold per LN slice (n = 5 slices, LN slices obtained from 3 mice). Two-way ANOVA, followed by Sidak posthoc test. **p < 0.01.

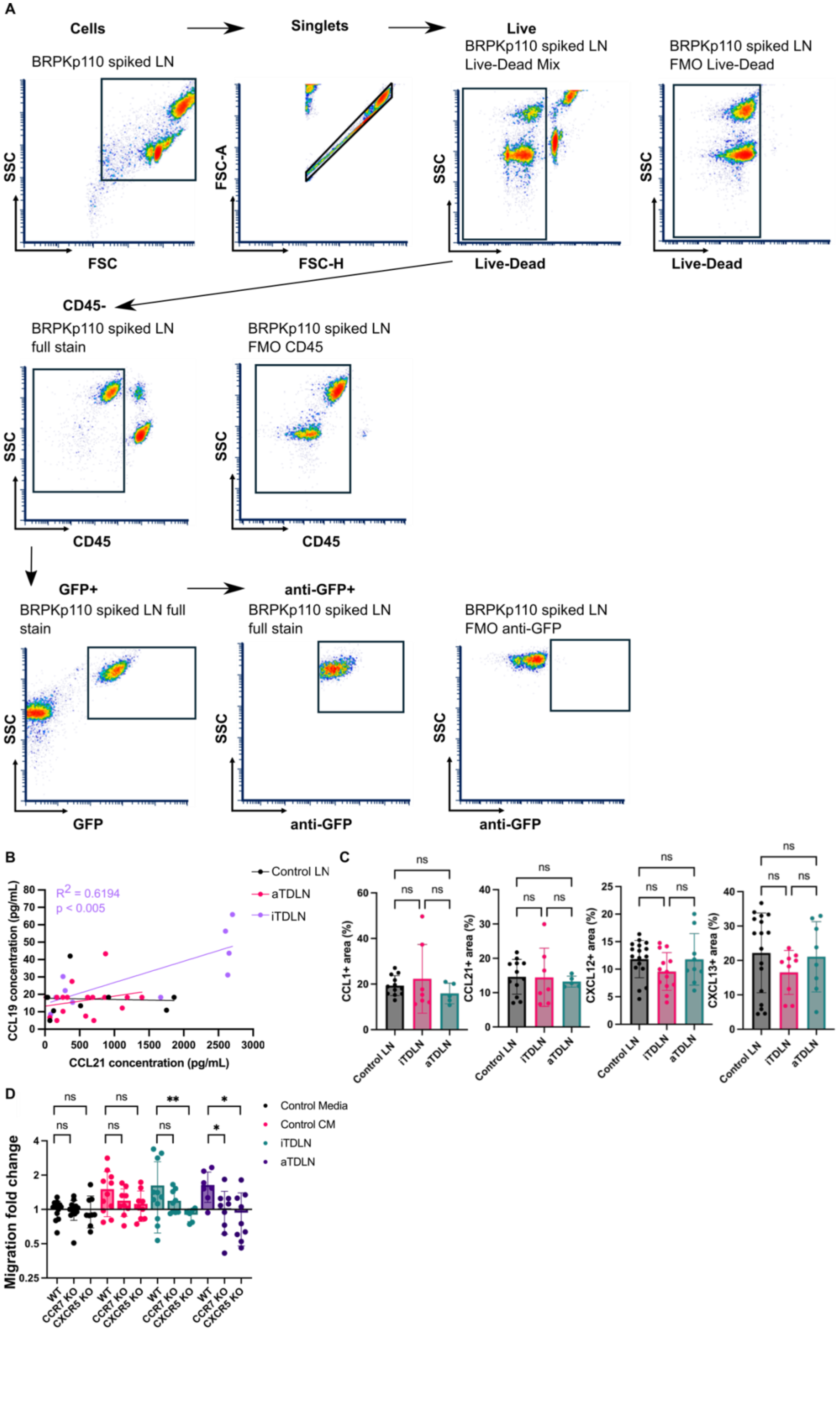
A
Cells
Singlets
Live
BRPKp110 spiked LN
BRPKp110 spiked LN
Live-Dead Mix
BRPKp110 spiked LN
FMO Live-Dead
SSC
FSC
FSC-A
FSC-H
Live-Dead
CD45-
BRPKp110 spiked LN
full stain
BRPKp110 spiked LN
FMO CD45
CD45
GFP+
anti-GFP+
BRPKp110 spiked LN full stain
BRPKp110 spiked LN
full stain
BRPKp110 spiked LN
FMO anti-GFP
GFP
anti-GFP
B
Control LN
aTDLN
iTDLN
$R^2$ = 0.6194
p < 0.005
CCL19 concentration (pg/mL)
CCL21 concentration (pg/mL)
C
CCL1+ area (%)
CCL21+ area (%)
CXCL12+ area (%)
CXCL13+ area (%)
ns
Control LN
iTDLN
aTDLN
D
Control Media
Control CM
iTDLN
aTDLN
Migration fold change
WT
CCR7 KO
CXCR5 KO
ns
**
*

**Figure S6: Quantification of BRPKp110 cell invasion in d5 TDLNs, characterization of immobilized chemokine in d5 TDLNs and its impact of BRPKp110 migration.** (A) Gating strategy for quantification of GFP+ BRPKp110 cancer cells in TDLNs. Suspension of cell from control LN was mixed with BRPKp110 cancer cells for a "spiked" cancer cell-positive control. (B) Correlation of CCL21 and CCL19 chemokine levels in CM obtained from control LN or from pre-metastatic iTDLN and aTDLN. Linear regression and correlation between CCL21 and CCL19 levels were done with paired data from supernatants obtained from culture of Control LNs, iTDLNs and aTDLNs. Each data point represents the supernatant from one LN slice (n=9-15/per group, LN slices obtained from 3 mice). Pearson $r = 0.7870$, R squared = 0.6194, **$p < 0.01$ for the iTDLN. (C) Fraction of total LN area positive for immobilized chemokines measured in control LN and pre-metastatic iTDLN and aTDLN after 1h of culture. Mean ± stdev; each data point represents a LN slice (n=5-8/per group, LN slices obtained from 3 mice). One-way ANOVA, followed by Tukey posthoc test. $p > 0.05$.(D) Migration of WT and KO BRPKp110 towards TDLN CM. Mean ± stdev; each data point represents the mean migration fold change per membrane, calculated from three non-overlapping fields of view (n = 3-4 membranes/condition; normalized data pooled from 3 independent in vitro experiments). Two-way ANOVA, followed by Tukey posthoc test. *$p < 0.05$, **$p < 0.01$.

**Table S1: Fluorescent antibodies used to label cells for flow cytometry and fluorescent microscopy.**

| Resource | Vendor | Identifier |
|---|---|---|
| anti-mouse CD16/32, clone 93, Rat IgG2a, κ | BioLegend | 101302 |
| anti-mouse Ki-67 antibody, clone 16A8, Isotype Rat IgG2a, κ | BioLegend | 652407 |
| anti-mouse APC CD197 (CCR7) clone 4B12, Rat IgG2a, κ | BioLegend | 120107 |
| anti-mouse APC CD198 (CCR8) clone SA214G2, Rat IgG2b, κ | BioLegend | 150309 |
| anti-mouse APC CD184 (CXCR4) clone L276F12, Rat IgG2b, κ | BioLegend | 146507 |
| anti-mouse APC CD185 (CXCR5) clone L138D7, Rat IgG2b, κ | BioLegend | 145505 |
| Isotype Control APC RatIgG2a, clone RTK2758, Rat IgG2a, κ | BioLegend | 400511 |
| Isotype Control APC Rat IgG2b, κ, clone RTK4530 | BioLegend | 400611 |
| Briliant Violet 421 anti-mouse podoplanin, clone 8.1.1, Syrian Hamster IgG | BioLegend | 127423 |
| Alexa Fluor 647 anti-mouse lyve1, clone ALY7, Rat IgG1,κ | Invitrogen | 53-0443-82 |
| Alexa Fluor 488 anti-mouse B220, clone RA3-6B2, Rat IgG2a, κ | BioLegend | 103229 |
| Starbright Violet 670 anti-mouse CD19, clone 6D5 , Rat IgG2a, κ | Bio-Rad | MCA1439SBV67 |
| PE anti-mouse CD45, clone 30-F11, Isotype Rat IgG2b, κ | BioLegend | 103105 |
| APC anti-GFP Antibody, clone FM264, Rat IgG2a, κ | BioLegend | 338010 |

**Table S2: Sequences of crRNA and protospacer-adjacent motifs used to generate BRPKp110 KO cell lines.**

| crRNA | Sequence (5′-3′) | PAM |
|---|---|---|
| CCR7-1 | CATCGGCGAGAATACCACGG | TGG |
| CCR7-2 | GTACAGGGTGTAGTCCACCG | TGG |
| CCR7-3 | CCTGGACGATGGCTACGTAG | CGG |
| CCR8-1 | TCGTGGGCTGCAAGAAACTG | AGG |
| CCR8-2 | CCTTGATGGCATAGACAGCG | TGG |
| CCR8-3 | TCTTGGATGGATGTGCCACG | AGG |
| CXCR5-1 | TTGGTGCGTAGAATCCACGA | GGG |
| CXCR5-2 | GTGGATTCTACGCACCAATG | GGG |
| CXCR5-3 | TACCCACTAACCCTGGACAT | GGG |
